# Unsupervised characterization of dynamic functional connectivity reveals age-associated differences in temporal stability and connectivity states during rest and task

**DOI:** 10.1101/2021.07.08.451590

**Authors:** Nisha Chetana Sastry, Dipanjan Roy, Arpan Banerjee

## Abstract

Understanding brain functions as an outcome of underlying neuro-cognitive network mechanisms in rest and task requires accurate spatiotemporal characterization of the relevant functional brain networks. Recent endeavours of the Neuroimaging community to develop the notion of dynamic functional connectivity is a step in this direction. A key goal is to detect what are the important events in time that delimits how one functional brain network defined by known patterns of correlated brain activity transitions into a “new” network. Such characterization can also lead to more accurate conceptual realization of brain states, thereby, defined in terms of time-resolved correlations. Nonetheless, identifying the canonical temporal window over which dynamic functional connectivity is operational is currently based on an ad-hoc selection of sliding windows that can certainly lead to spurious results. Here, we introduce a data-driven unsupervised approach to characterize the high dimensional dynamic functional connectivity into dynamics of lower dimensional patterns. The whole-brain dynamic functional connectivity states bearing functional significance for task or rest can be explored through the temporal correlations, both short and long range. The present study investigates the stability of such short- and long-range temporal correlations to explore the dynamic network mechanisms across resting state, movie viewing and sensorimotor action tasks requiring varied degrees of attention. As an outcome of applying our methods to the fMRI data of a healthy ageing cohort we could quantify whole-brain temporal dynamics which indicates naturalistic movie watching task is closer to resting state than the sensorimotor task. Our analysis also revealed an overall trend of highest short range temporal network stability in the sensorimotor task, followed by naturalistic movie watching task and resting state that remains similar in both young and old adults. However, the stability of neurocognitive networks in the resting state in young adults is higher than their older counterparts. Thus, healthy ageing related differences in quantification of network stability along task and rest provides a blueprint of how our approach can be used for cohort studies of mental health and neurological disorders.

## 1. Introduction

Functional connectivity (FC) - most simplistically computed using the pairwise Pearson correlations between brain regions using blood oxygen level dependent (BOLD) fMRI has proven to be a powerful tool for studying the functional organization of the brain (Friston, et al. 1993). FC sheds light on the functional coupling and connectedness between proximal and distal brain regions subserving crucial role towards the neuronal processing of a task (Aertsen, Gerstein, Habib, & Palm, 1989) (Friston, Frith, Liddle, & Frackowiak, 1993). However, emergence of superior computing prowess has allowed us to critique the inferences drawn from time-averaged, static FC usually computed by collapsing the functional dynamics across time (Ciric, Nomi, Uddin, & Satpute, 2017) (Mash, et al., 2019). Recently, dynamic functional connectivity (dFC) has emerged as a major topic in the resting-state BOLD fMRI literature (Hutchison et al., 2013). The more refined measure of Dynamic functional connectivity (dFC) is commonly computed using the sliding window framework, which estimates dFC by computing average FC over small windows of time, and subsequently sliding the window over the entire duration of the BOLD time series (Hutchison & et al, 2013). Although, the sliding window approach has been the most common, simple, and intuitive analysis strategy for estimating dFC (Kudela, Harezlak, & Lindquist, 2017) (Preti, Bolton, & Van De Ville, 2017), the method suffers from prominent drawbacks. Arbitrary choice of window length, inherent variation present in the estimate that can be confused with the empirical time-varying nature of FC, equal weighting of all observations within the window leading to spurious fluctuations being magnified – all add to the woes of sliding window based approach (Lindquist, Xu, Nebel, & Caffo, 2014) (Hindriks, et al., 2016) (Preti, Bolton, & Van De Ville, 2017).Over the years, many meaningful extensions have been suggested to improve sliding window approach. Independent component analysis (ICA) was used to decompose windowed BOLD timecourses (Kiviniemi, et al., 2011). Several graph theoretical summary measures such as assortivity, modularity, efficiency offer promising avenues to extract information from dFC (Bullmore & Sporns, 2009). In addition, clustering algorithms such as K- means clustering (Damaraju, et al., 2014) (Allen, et al., 2014), hidden Markov models (HMM) (Vidaurre, Smith, & Woolrich, 2017), temporal ICA (TICA) (Yaesoubi, Miller, & Calhoun, 2015) allows to identify clustering-derived recurring connectivity patterns or dFC states. Several conceptual alternative strategies such as wavelet transform coherence (Chang & Glover, 2010), a time/frequency analysis strategy with an observation window for the frequency content of the time courses; and frame-wise analysis of the BOLD timecourses (Cabral, et al., 2017), which allows information to be retrieved from the observation of single frames and yield temporally subsequent co-activation maps (Liu, Chang, & Duyn, 2013); have been suggested (see (Preti, Bolton, & Van De Ville, 2017) for a review).

In spite of inherent limitations, dFC captures the fluctuations of FC, which contain meaningful information on minute temporal scale (Hutchison & et al, 2013). While, accounting these fluctuations are critical for understanding complex behaviour, nonetheless, stable representation of information of neural activity and corresponding stability of FC patterns over time is critical for survival (Li, Lu, & Yan, 2019). In other words, in an axiomatic sense, there must exist a temporally stable connectivity pattern that corresponds to one or multiple functional states of the brain. Subsequent transition between successive functional brain states can be characterized by estimating the dFC patterns. The two most widely applied dynamic measures in brain/behaviour analyses that are constructed from dFC time courses based on either applying sliding window or frame wise analysis are connection variability (CV) and connectivity states (CS). Stable representation of information processing will reflect the robustness of recurring patterns of CS concatenated across subjects to influence myriad of behaviour (Bolton, Morgenroth, Preti, & Van De Ville, 2020). Identifying both connectivity states and their stability in brain dynamics requires delimiting the dFC evolution and dFC stability with measures of optimality which may or may not be crucially linked with the underlying subject specific structural connectivity (Surampudi, et al., 2019). The goal of this article is to develop an unsupervised approach to characterize optimal dFC states and go beyond what has been proposed in the existing literature based on widely applied sliding window or frame wise based approach (Bolton, Morgenroth, Preti, & Van De Ville, 2020).

Previous studies exploring temporal dynamics of FC have tried to investigate the stability by calculating the correlation between FC matrices computed from successive temporal windows (Hansen, Battaglia, Spiegler, Deco, & Jirsa, 2015), characterizing CV of the functional connectivity profile of a given region across time (Zhang & et al, 2016) (Guo, Zhao, Tao, Liu, & Palaniyappan, 2017), by estimating voxel level dFC maps using Kendall’s coefficient of concordance with time windows as raters (Li, Lu, & Yan, 2019), by estimating the standard deviation of global modularity averaged across all timepoints and all participants (Hilger, Fukushima, Sporns, & Fiebach, 2019). FC stability has been shown to increase with motor learning (Yu, Song, Huang, Song, & Liu, 2020), decrease in patients of schizophrenia and their siblings (Guo, Zhao, Tao, Liu, & Palaniyappan, 2017), was significantly higher in patients with major depressive disorder (Demirtas, et al., 2016). These studies emphasise the neurobiological significance of the stability of FC.

Quantifying the temporal stability of dFC patterns is of immediate concern to studies investigating the relationship between resting state and task-related brain dynamics (Li, Lu, & Yan, 2019). Spontaneous brain activity during rest is not random and show specific spatio-temporal organization in state space (Deco, Jirsa, & Mcintosh, 2011). From a dynamical systems point of view, the resultant emerging resting state functional connectivity of the brain networks, quantitatively, fits best with the experimentally observed functional connectivity when the brain network operates at the edge of instability. Under these near critical conditions, the slow fluctuating (< 0.1 Hz) resting state BOLD networks emerge as structured noise fluctuations around a stable low firing activity equilibrium state in the presence of latent "ghost" multi stable attractors (Deco & Jirsa, 2012). Recent work has further demonstrated that during spontaneous resting state activity the ghost attractors makes frequent excursion to functionally and behavioural relevant phase locking states in a low dimensional state space (Vohryzek, Deco, Cessac, Kringelbach, & Cabral, 2020) . Brain resides in a specific attractor state defined by a certain FC pattern according to the cognitive demands of the task (Fedorenko & Thompson-Schill, 2014) (Pillai & Jirsa, 2017). An overall increase in FC stability has been reported in the presence of the task (Gonzalez-Castillo & Bandettini, 2018). Subsequently, temporal stability of FC guides the stability of a functional state. Thus, we tested the following hypothesis, unsupervised dFC characterization will reveal task specific dFC stability patterns that are local in time, whereas for the resting state dFC patterns, these functional states are composed of non-local correlations in time. Prior studies have also explored changes in temporal stability of functional architecture in resting state of healthy control and patients with psychiatric disorders, and different battery of tasks. Zhang and colleagues showed disorder specific (ADHD, schizophrenia, autism spectrum disorder) variability modifications in functional architecture of DMN, visual and subcortical regions of the brain (Zhang & et al, 2016). Increased functional stability in high-order visual regions during naturalistic movie watching task were identified (Li, Lu, & Yan, 2019), but these studies are limited to stability of FC of a given region. The second test for an unsupervised dFC characterization algorithm will be application to a specific neuroscience problem, such as investigation of lifespan trajectories in healthy ageing. Although previous studies have explored the association between dynamic functional connectivity and age (Viviano, Raz, Yuan, & Damoiseaux, 2017) (Chen, et al., 2017) (Xia, et al., 2018), how the stability of functional architecture modifies across age remains an open question.

The aim of the present study is three-fold: 1) to precisely characterise the stability of whole-brain dFC patterns 2) to demonstrate that dFC patterns are locally stable during task 3) identify the dFC patterns during task and rest for a cross-sectional population with age range over human adult lifespan (18-88 years). This manuscript is organized as follows. First, we estimate BOLD phase coherence over time (Glerean, Salmi, Lahnakoski, Jääskeläinen, & Sams, 2012) which was used as a measure of dFC for rest and task. Next, we proceed with unsupervised characterization of dFC subspaces involved in task and rest. Subsequently, the temporal stability of dFC subspaces were computed using two different measures - angular separation and the Mahalanobis distance (Mahalanobis, 1930) (Shen, Kim, & Wang, 2010). Finally, the temporal stability of dFC was analysed to draw critical insights about age associated differences to task and rest using a large human cohort of healthy ageing (Shafto, et al., 2014).

## 2. Methods

### 2.1 Data sources and participants

The data were collected as part of stage 2 of the Cambridge Centre for Ageing and Neuroscience (CamCAN) project (available at http://www.mrc-cbu.cam.ac.uk/datasets/camcan) (Taylor, et al., 2017) (Shafto, et al., 2014). The CamCAN is a large-scale multimodal, cross-sectional, population- based study. The database includes raw and pre-processed structural magnetic resonance imaging (MRI), resting state and active tasks using functional MRI (fMRI) and Magnetoencephalogram (MEG), behavioural scores, demographic and neuropsychological data. From 3000 participants of stage 1, a subset of approximately 700 participants who were cognitively healthy (MMSE score >25), with no past or current treatment for drug abuse or usage, met hearing threshold greater than 35 dB at 1000 Hz in both ears, had at least a corrected near vision of 20/100 with both eyes and could speak English language (native English speaker or bilingual English from birth) were eligible for MRI scanning. They were home interviewed and recruited to stage 2. The study was in compliance with the Helsinki Declaration and was approved by the Cambridgeshire 2 Research Ethics Committee. The fMRI data from resting state and task periods (naturalistic movie watching and sensorimotor task) was used in the present study.

### 2.2 Data acquisition and experimental paradigm

The fMRI data were collected at MRC Cognition and Brain Sciences Unit, on a 3T Siemens TIM Trio scanner with a 32-channel head coil, the head movement was restricted with the aid of memory foam cushions. For the tasks, the instructions and visual stimuli were back projected onto the screen, auditory stimuli were presented via MR-compatible Etymotics headphones and manual responses from the participants made with the right hand were recorded using an MR-compatible button box (Taylor, et al., 2017). The fMRI data for eyes-closed resting state and sensorimotor task were acquired using Echo-Planar Imaging (EPI) sequence, consisted of 261 volumes, each volume with 32 axial slices (slice thickness 3.7mm, interslice gap 20%) acquired in descending order, TR 1970 ms, TE 30 ms, voxel-size 3 mm 3 mm 4.44 mm. The duration of both the scans was 8 min 40s. The fMRI data for the naturalistic movie watching task were acquired using multi-echo EPI sequence, consisting of 193 volumes of 32 axial slices each (slice thickness 3.7mm, interslice gap 20%) acquired in descending order, TR 2470 ms, TE [9,4,21.2,33,45,57] ms, voxel-size 3 mm 3 mm 4.44 mm. The duration of the scan was 8 min 13s.

The task-induced BOLD data from the naturalistic movie watching task was acquired from participants, who watched 8 minutes of narrative preserved, condensed, black and white version of Alfred Hitchcock’s television drama “Bang! You’re Dead”. The participants were not aware of the title of the movie but were instructed to pay attention to the movie. In the sensorimotor task, the trials consisted of a binaural tone simulation at either 300, 600, or 1200 Hz and bilateral black and white checkerboard. The participants were asked to button press with their right index finger if they hear or see any stimuli. More details about the task paradigm have been presented here (Shafto, et al., 2014) (Taylor, et al., 2017).

### 2.3 Data pre-processing

Pre-processed data was provided by Cam-CAN research consortium. Mean regional BOLD time series were estimated in 116 parcellated brain areas of Anatomical Automatic Labelling atlas (AAL) (Tzourio-Mazoyer, et al., 2002) (available at http://www.gin.cnrs.fr/tools/aal). We selected 50 participants, 25 were young adults (48% female; mean age = 24.1 士 3.33 years) randomly selected from age range 18-28, and remaining 25 were old adults (52% female; mean age = 63.8士2.63 years) randomly selected over age range 60-68 years. Each participant’s BOLD time series in the resting state, naturalistic movie watching and sensorimotor tasks were extracted.

### 2.4 Data analysis

#### 2.4.1 Characterization of dynamic functional connectivity

Time-resolved dynamic functional connectivity (dFC) was estimated, for each individual, using BOLD phase coherence (**Figure 1A)** (Glerean, Salmi, Lahnakoski, Jääskeläinen, & Sams, 2012) (Ponce-Alvarez, et al., 2015) (Deco & Kringelbach, 2016) (Cabral, et al., 2017), which resulted in a matrix with size *NxNxT*, where N=116 is the number of brain regions defined by AAL atlas, T is the total number of time points (T=261 for resting state and Sensorimotor task, T=193 for naturalistic movie watching task). We chose BOLD phase coherence instead of computing correlation over a sliding window to calculate dFC, because BOLD phase coherence is an instantaneous measure with maximum temporal resolution (Glerean, Salmi, Lahnakoski, Jääskeläinen, & Sams, 2012). BOLD phase coherence does not require time-windowed averaging, that generates biased estimates if the window length is short and reduces temporal resolution if the window length is longer (Glerean, Salmi, Lahnakoski, Jääskeläinen, & Sams, 2012).

**Figure 1:**
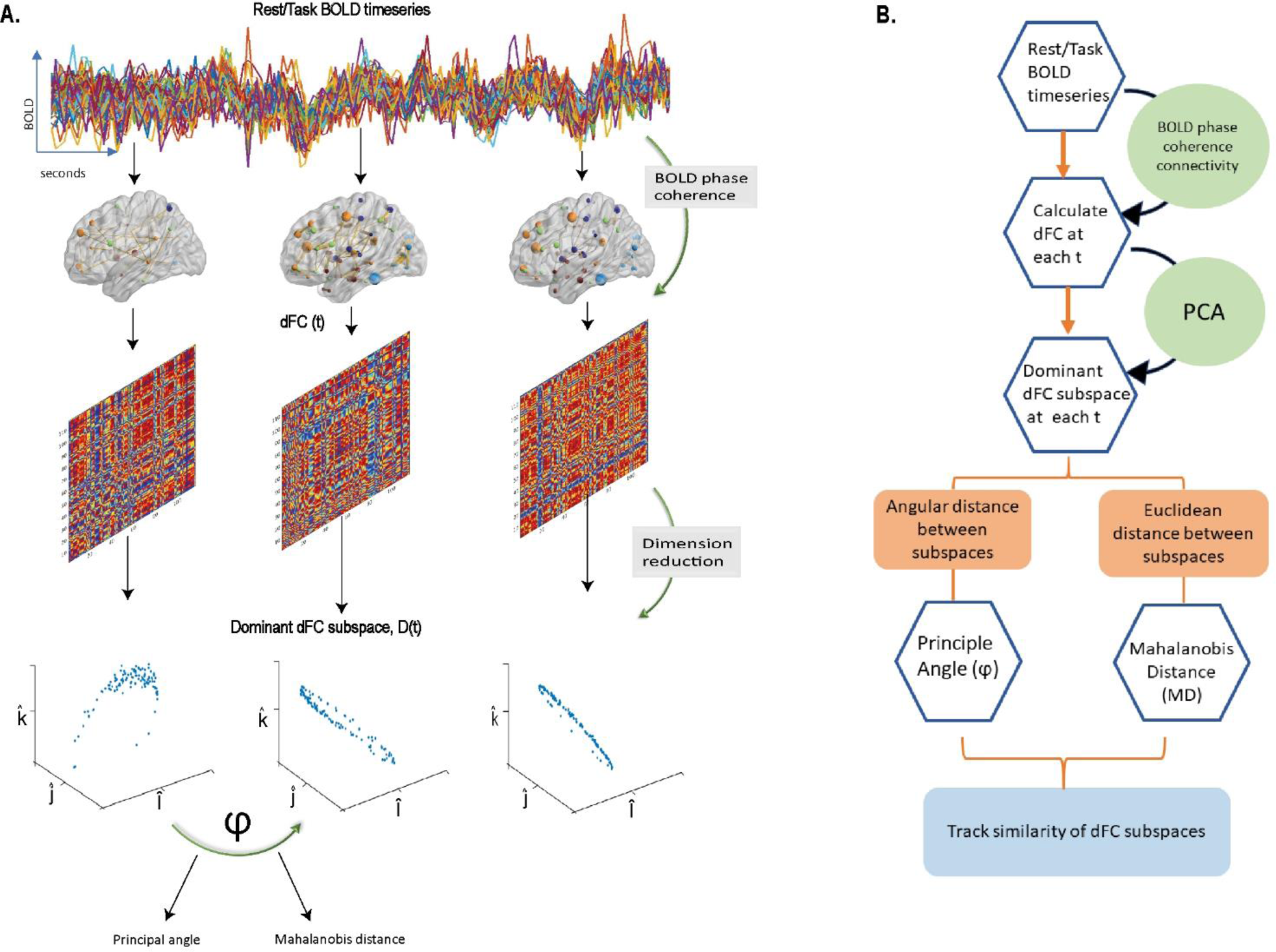
**(A)**. The schematic diagram shows how the temporal stability of dynamic functional connectivity subspaces (dFC) are computed. Dominant dFC subspace, at each time point, is estimated using the first three principal components of dFC(t), that was computed using the measure of BOLD phase coherence. The similarity between dFC subspaces are calculated using principal angle (Angular distance) and Mahalanobis distance (Euclidean distance). If the dominant dFC subspaces are similar for extended timepoints, then they are considered to be stable. **(B).** A flowchart representation of the method

First, the instantaneous phases *θ*(*n*, *t*) of the BOLD time series for all the brain regions,*n*, was computed using Hilbert transform. The real-valued modulated BOLD signal *s*(*t*) is expressed as an analytical signal in the complex plane as:

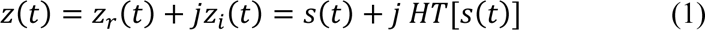

Where, (HT [*]) represents the Hilbert transform. The instantaneous phase *θ* (*t*) is computed as follows:

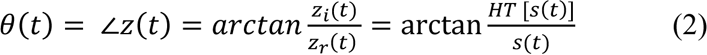

Given the phases of the BOLD time series, phase coherence i.e., *dFC* (*n*, *p*, *t*) for brain regions, *n* and *p* at time *t* is computed as:

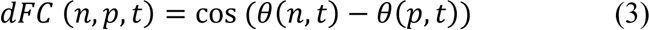

when, the phases of BOLD signals, *θ*(*n*, *t*), *θ*(*p*, *t*) of the brain regions *n*, *p* are synchronized, *dFC*(*n*, *p*, *t*) (ranges from -1 to 1) is close 1, when the phases from the BOLD signals of brain regions *n*, *p* are orthogonal *dFC*(*n*, *p*, *t*) is close to 0. Since the phases are undirected, *dFC*(*n*, *p*, *t*) is symmetric along the diagonal.

In addition to this, to check for reliability, we compute dFC using a sliding-window approach (Hutchison & et al, 2013) with non-overlapping, gaussian windows, varying the window length (10, 20, 30 time points) (Supplementary information - **S 1**, **S 2**, **S 3**).

#### 2.4.2 Extracting Dominant dynamic functional connectivity

Principal component analysis (PCA) was applied to participant-wise *dFC* (*n*, *p*, *t*) matrix of size *NXN* representing the FC between *n*th and *p*th brain area for each time point. PCA is an unsupervised, multivariate dimension reduction method that decomposes the data into a set of orthogonal principal components or leading eigenvectors sorted by their contribution to the overall variance (Friston, 1993). Thus, *dFC* (*n*, *p*, *t*) or simply *dFC*_*t*_ can be expressed as

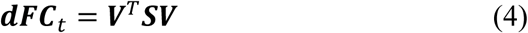

where, matrix *V* of size *NXN* are set of eigenvectors, with each column of *V* of size 1*XN* representing orthogonal principal component, and *S* the diagonal matrix 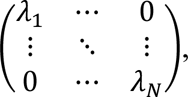 such that *λ*_1_ > *λ*_2_ … > *λ*_*N*_

If *k* is the number of principal components chosen to represent *dFC*, the corresponding subspace *D*(*n*, *k*, *t*) or *D*_*t*_, representative of dominant dFC pattern, can be expressed as

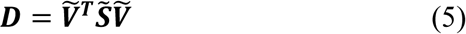

where, *Ṽ*^*T*^ is a dimensionally reduced matrix of size *N X k S̃* is a diagonal matrix 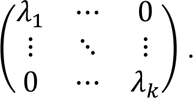

In this study, we chose k = 3 because for all participants at least 99% variance in *dFC* matrix is captured by the 3 leading eigenvectors (**S 4**). The dimension of *dFC*(*n*, *p*, *t*) has been reduced to *D*(*n*, *k*, *t*).

#### 2.4.3 Computation of stability of dynamic functional architecture

We seek to characterize the temporal stability of the dominant subspace *D*(*n*, *p*, *t*) (or referred to as simply *D*_*t*_) by estimating how similar they are across time *t*. To estimate the similarity between dominant dFC configurations, we introduce two types of distance measures successive dFC subspaces, 1) angular distance 2) Normalised Euclidean distance **(Figure 1B).** We define angular distance as the principal angle between the dFC subspaces from different time points, given by the following equation:

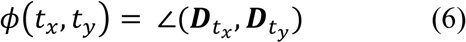

Where, each entry in the time X time *temporal stability matrix*, *ϕ*(*t*_*x*_, *t*_*y*_) is the principal angle between the two N X k dimensional subspaces at *t*_*x*_ and *t*_*y*_ (Banerjee, Pillai, Sperling, Smith, & Horwitz, 2012) ( Björck & Golub, 1973). The principal angle ranges between 0 (low angular distance) to π/2 (high angular distance).

For each individual, we calculate the angular distance between dominant dFC subspaces at *t*_*x*_ and *t*_*y*_, by estimating the principal angle between them. The low principal angle between dominant dFC subspaces means that their dFC configurations are very similar. On the contrary, the high principal angle between dominant dFC subspaces means that their dFC configurations are dissimilar.

We define the normalised Euclidean distance between dominant dFC subspaces by the Mahalanobis distance. Mahalanobis distance measures the distance between points in space 1 from space 2 with the following equation:

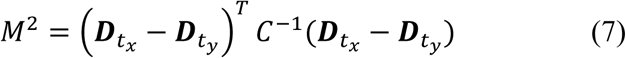

where *M*^2^ is the distance between each entry of *D*_*t*_ and *D*_*t*_ Subsequently, for each individual, we estimate the time X time *temporal stability matrix*, where each entry is the Mahalanobis distance (*M* ranges between 0.5 to 2.5), averaged across all brain parcels. Low *M* means that dominant dFC subspaces are similar, high *M* means that the dFC subspaces are dissimilar.

#### 2.4.4 Quantifying complexity of temporal stability matrices

*Entropy:*

To evaluate the informational content of temporal stability matrices we evaluated the entropy, for all three categories, rest, movie viewing and sensorimotor task in young and old adults. Entropy is defined by the following equation:

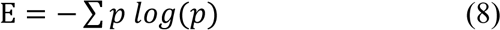

where *p* contains the normalised histogram counts returned from ‘imhist.m’. ‘imhist.m’ calculates the histogram of temporal stability matrices and returns histogram counts.

Entropy in temporal stability matrices for empirical data and surrogate data were compared by generating random time series using MATLAB function ‘randn.m’ and down sampling them at 0.1 Hz mimicking BOLD activity of each subject. The *D* matrices with same dimensions as the empirical data were computed. Welch’s corrected t-tests revealed significant differences between the entropy of surrogate and empirical temporal stability matrices of rest (p=0.000464) (**S 5**).

*Frobenius norm:*

Frobenius norm was used to measure the differences between the temporal stability matrices computed for rest and the task conditions. Frobenius norm, also called the Euclidean norm of a matrix, is defined as the square root of the sum of the absolute squares of its elements. Here, we calculate Frobenius norm between temporal stability matrices with the following equation:

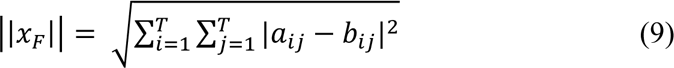

where *a*_*ij*_ and *b*_*ij*_ are the entries in the temporal dynamic matrices of rest and any of the task conditions respectively (movie watching or sensorimotor). *x*_*F*_ is also computed between the two tasks.

*Stochastic characterization of dFC*

The temporal variation of two measures, principal angle and Mahalanobis distance between the dominant *dFC* subspaces essentially capture the degree of temporal variation in functional network. Principal angular values close to 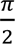 or high Mahalanobis distance at a specific time point reflects the reorganization of the functional state itself, whereas angular values closer to zero or low Mahalanobis distance indicates minor deviation from previous time. To understand the underlying stochastic characteristics of these measures, we use auto-regressive (AR) models where present values of *ϕ*(*t*)/ *ϕ*(*t*) are modelled as a linear weighted sum of values from past *ϕ*(*t* − 1), *ϕ*(*t* − 2) … *ϕ*(*t* − *i*)/*ϕ*(*t* − 1), *ϕ*(*t* − 2) … *ϕ*(*t* − *i*). The AR (*ρ*) process, *X*_*t*_ (*ϕ*(*t*) or *ϕ*(*t*)) is given by the following equation:

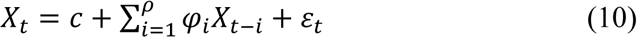

where *φ*_1_ … … … … *φ*_*ρ*_ are parameters of the model, *c* is a constant, *ε*_*t*_ is white noise and *ρ* is the lag term or model order. The simplest AR process is AR (0) is essentially a white noise process. In AR (1), the current value is dependant only on its immediately preceding value, and hence captures a Markovian process. Optimal model of an AR process can be computed using the Akaike information criterion (AIC) which is expressed as

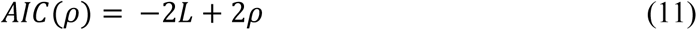

where *L* is the likelihood function computed by summing up over the mean squared error for an AR model of order *ρ* (Wagenmaker & Farrell, 2004) (H.Akaike, 1974).Optimal model order can be selected at a value of *ρ* where AIC is minimum. We varied the model order (*ρ*) from 0 to 100 and use the first minimal AIC value to select the best AR (*ρ*), model. If the model order is found to be greater than 1, the underlying process is considered non-Markovian.

## 3 Results

### 3.1 Dynamic functional connectivity (*dFC*) patterns during rest, continuous naturalistic movie watching, and discrete sensorimotor task

We computed the *dFC* from parcellated BOLD time series of resting state, naturalistic movie watching task where the participants watched and listened to an excerpt from Alfred Hitchcock’s “Bang! You’re Dead”, and a sensorimotor task where participants responded by a button press to either a visual or an auditory stimulus from the Cam-CAN dataset (details in Methods). **Figure 2A** represents dFC obtained using BOLD phase coherence connectivity in resting state. We report the results of the analysis on young adults (age range 18-28) in this section.

**Figure 2:**
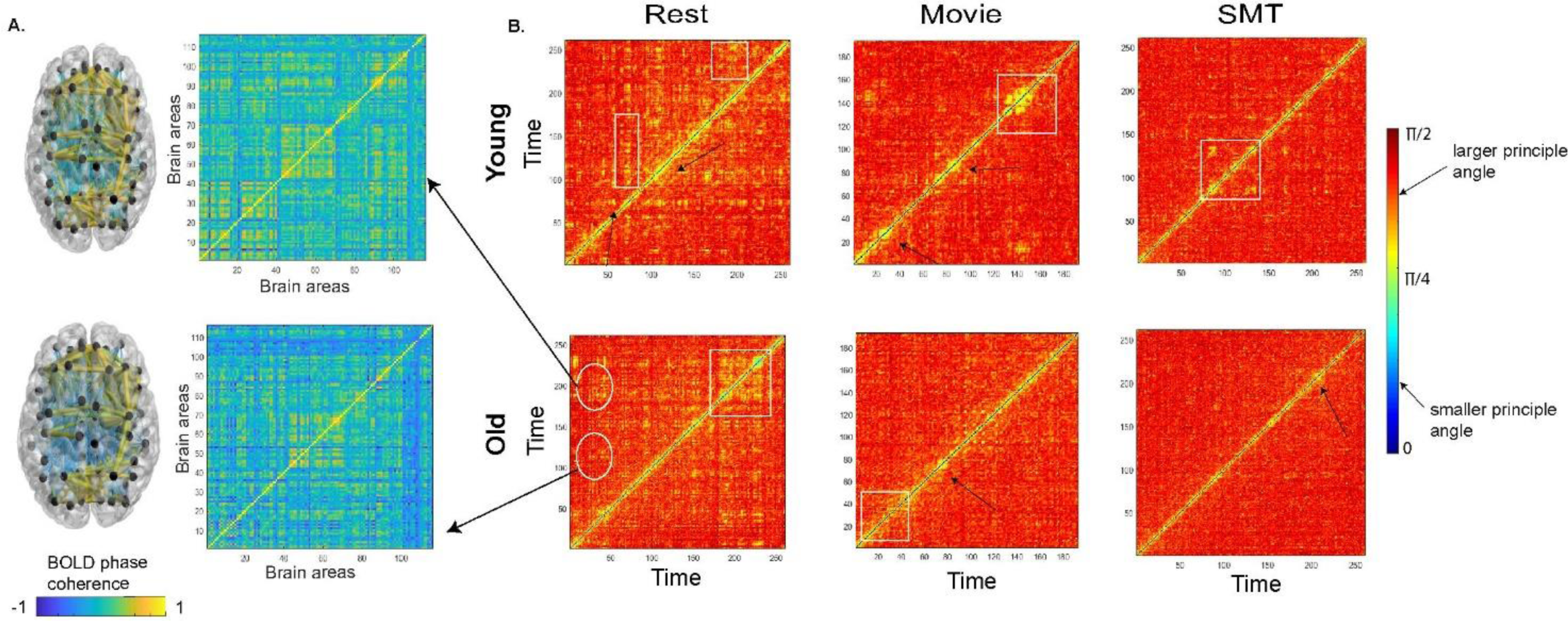
**(A)** dFC matrices estimated using BOLD phase coherence. **(B)** Time X Time temporal stability matrix of resting state, naturalistic movie watching task and discrete, sensorimotor task for young and old adults. Each entry in the matrix is the principal angle *ϕ*(*t*_*x*_, *t*_*y*_) between dominant dFC subspaces at *t*_*x*_ and *t*_*y*_. The principal angle ranges between 0 (low angular distance) to π/2 (high angular distance). Resting state, in both young and old adults, has shorter-lived, global spread of patterns of temporal stability. On the contrary, both the tasks have a longer-lived, local spread of patterns of stability (indicated by arrows and rectangular boxes).

Dominant *dFC* subspaces were obtained by applying the unsupervised approach of Principal Component Analysis (PCA) to BOLD time series at each time point, and then reconstructing either the task or rest as the dynamics of a reduced dimensional *dFC* subspace. To demonstrate, that the unsupervised characterization of *dFC* patterns indeed capture the functional brain network organization, we computed the differences between the temporal stability matrices of rest and the two task conditions; first using the measure of principal angles and second using the measure of Mahalanobis distance. Thereafter, other measures of complexity and temporal variability were tested.

#### 3.1.1 Using angular distance to characterize temporal stability matrices

First, we calculate the principal angles among the dominant *dFC* subspaces generated across all time points. This resulted in time X time temporal stability matrix, averaged across all subjects, where each entry in the matrix is the angle between dominant *dFC* subspaces at *t*_*x*_ and *t*_*y*_, as shown in **Figure 2BFigure 2**. We consider a dominant *dFC* configuration to be stable if the subsequent subspaces are similar in configuration, i.e., less “angular distant” for extended duration of time points. Results shown in **Figure 2B** indicate that the resting state has a global spread of shorter-lived, repeated patterns of stability than both tasks. On the contrary, both the task cohorts, passive movie watching, and sensorimotor task, showed a local spread of, longer-lived stability patterns suggesting that local temporal stability of functionally connected networks are higher in the task than in resting state. To quantify these observations, we calculated the entropy of temporal stability matrices of each category. The plots in **Figure 4A**, which represent entropy of temporal dynamic matrices of three categories, report resting state to have the highest entropy, followed by movie watching task and sensorimotor task. The distribution was parametric (normality check was done with Jarque-Bera test and verified with D’Agostino-Pearson omnibus test), paired two-sample t-tests and effect size analysis using Cohen’s d, revealed significant differences (at 95% significance level) in entropy values between resting state and movie watching task (p=0.0026 d=0.8), and resting state and sensorimotor task (p=0.001, d=1.009). However, difference in movie watching task and sensorimotor task were not significant (p=0.4907 ns). Further, to analyse how similar temporal stability matrices across rest and tasks are, we calculate the Frobenius norm as shown in **Figure 4B**. The results reveal a shorter Frobenius norm between the temporal dynamic matrices of the resting state and movie watching task, than the resting state and sensorimotor task.

#### 3.1.2 Using Mahalanobis distance to characterize temporal stability matrices

Alternatively, we evaluate the temporal stability of *dFC*, by estimating Mahalanobis distance, that resulted in a time X time temporal stability matrix. Each entry of this matrix is the Mahalanobis distance between dominant dFC subspaces (**Figure 3A)**. Results, as shown in **Figure 3B** and **Figure 3C**, reveal global, shorter-lived repeated patterns of temporal stability in resting state and local, longer-lived temporal stability patterns in both the tasks. The entropy results (**Figure 4A**) reveal high entropy in the resting state, followed by movie watching task and sensorimotor task. The distribution was non-parametric (normality check was done with Jarque-Bera test and D’Agostino-Pearson omnibus test), we employed Wilcoxon matched paired test to compute statistical significance between the entropy of temporal stability matrices of each category, although the results did not reveal statistical significance, the trend in entropy is similar to the trend in angular distance metric. We repeated the Frobenius norm analysis, which produced similar results as the angular distance metric, as shown in **Figure 4B.**

**Figure 3:**
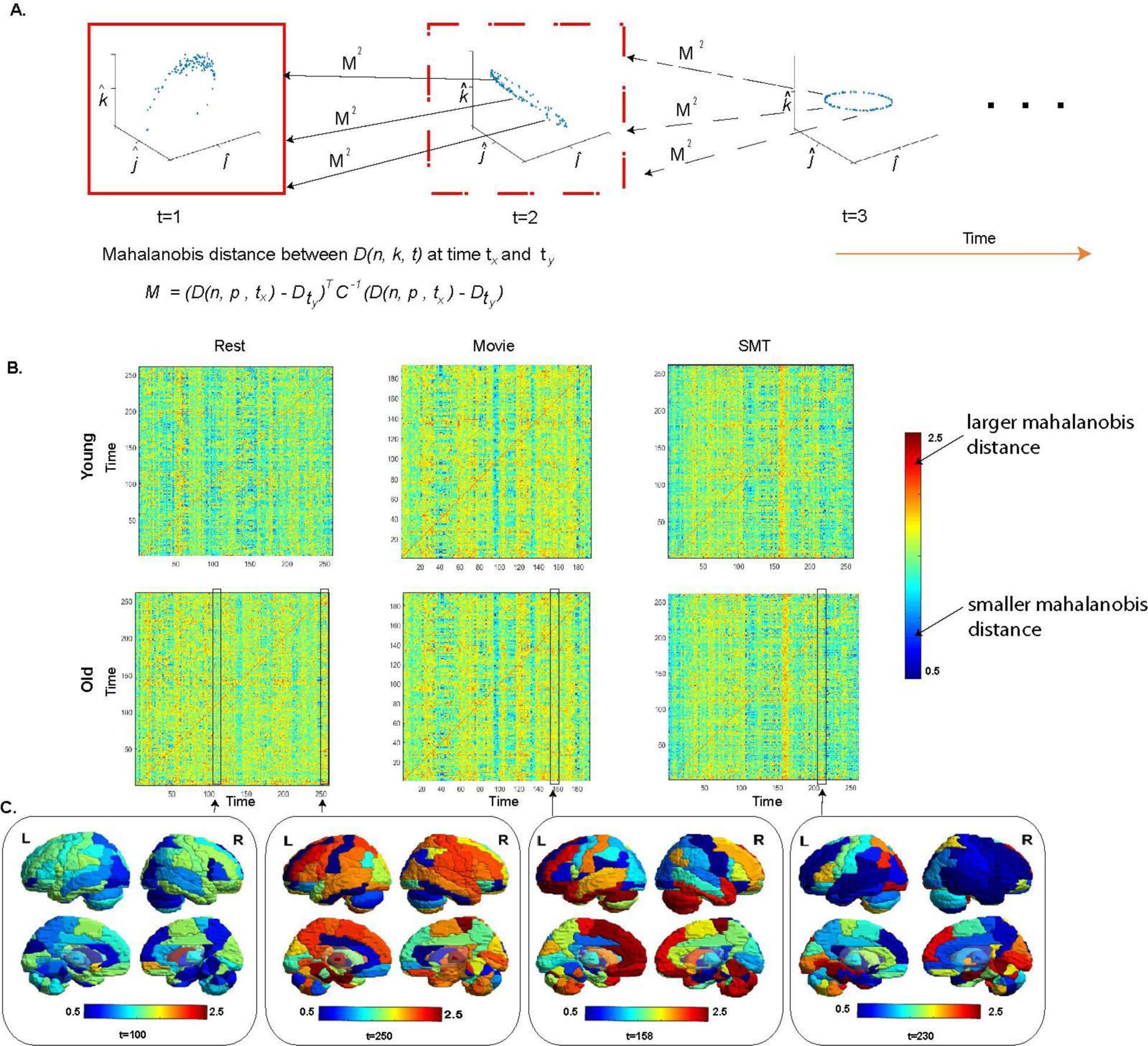
Mahalanobis distance between dominant dFC subspaces. **(A)** Mahalanobis distance is a pairwise Euclidean distance between the ROIs of dominant dFC subspace at *t*_*x*_, with the whole-brain dominant dFC subspace at *t*_*y*_. **(B)** Time X Time temporal stability matrix of resting state, naturalistic movie watching task, and sensorimotor task, where each entry in the matrix is Mahalanobis (*ϕ*^2^(*t*_*x*_, *t*_*y*_)) distance between the dominant dFC subspaces. Mahalanobis distance between dominant dFC subspaces is low when the dFC configurations are similar. **(C)** The profile of temporal stability estimated with Mahalanobis distance across the brain regions at different instances of time.

### 3.2 Unsupervised characterization of ***dFC*** across healthy ageing

Next, we have included two cohorts, young and old adults from the Cam-CAN dataset and carried out unsupervised characterisation of dFC using participant’s resting state, movie watching, and sensorimotor task data to identify age associated alterations in temporal stability of dominant *dFC* subspaces.

#### 3.2.1 Using principal angle to quantify temporal stability differences in dFC between young and elderly

The time X time temporal stability matrix was computed for the aged cohort (age range 60- 68) and compared with that of younger cohort computed in the section 3.1. A global spread of shorter duration of temporal stability patterns was observed in resting state and local spread of longer duration temporal stability patterns was observed in the task, in both young and old adults. Further, entropy analysis revealed **(Figure 4A)** a similar trend of peak entropy in resting state, followed by movie watching task and sensorimotor task in both young and old cohorts. The distribution was parametric (normality check was done with Jarque-Bera test and D’Agostino-Pearson omnibus test), paired two-sample t-test revealed significant differences in entropy values between resting state and movie watching task (p=0.000435, d=0.971), movie watching task, and sensorimotor task (p=0.0438, d=0.370), resting state and sensorimotor task (p=0.000567, d=1.319) of the older cohort (P values of young adults are reported in the previous section). The Frobenius norm analysis as shown in **(Figure 4B)** also revealed a similar trend in young and old adults i.e., shorter Frobenius norm between resting state and movie watching task than resting state and sensorimotor task

**Figure 4:**
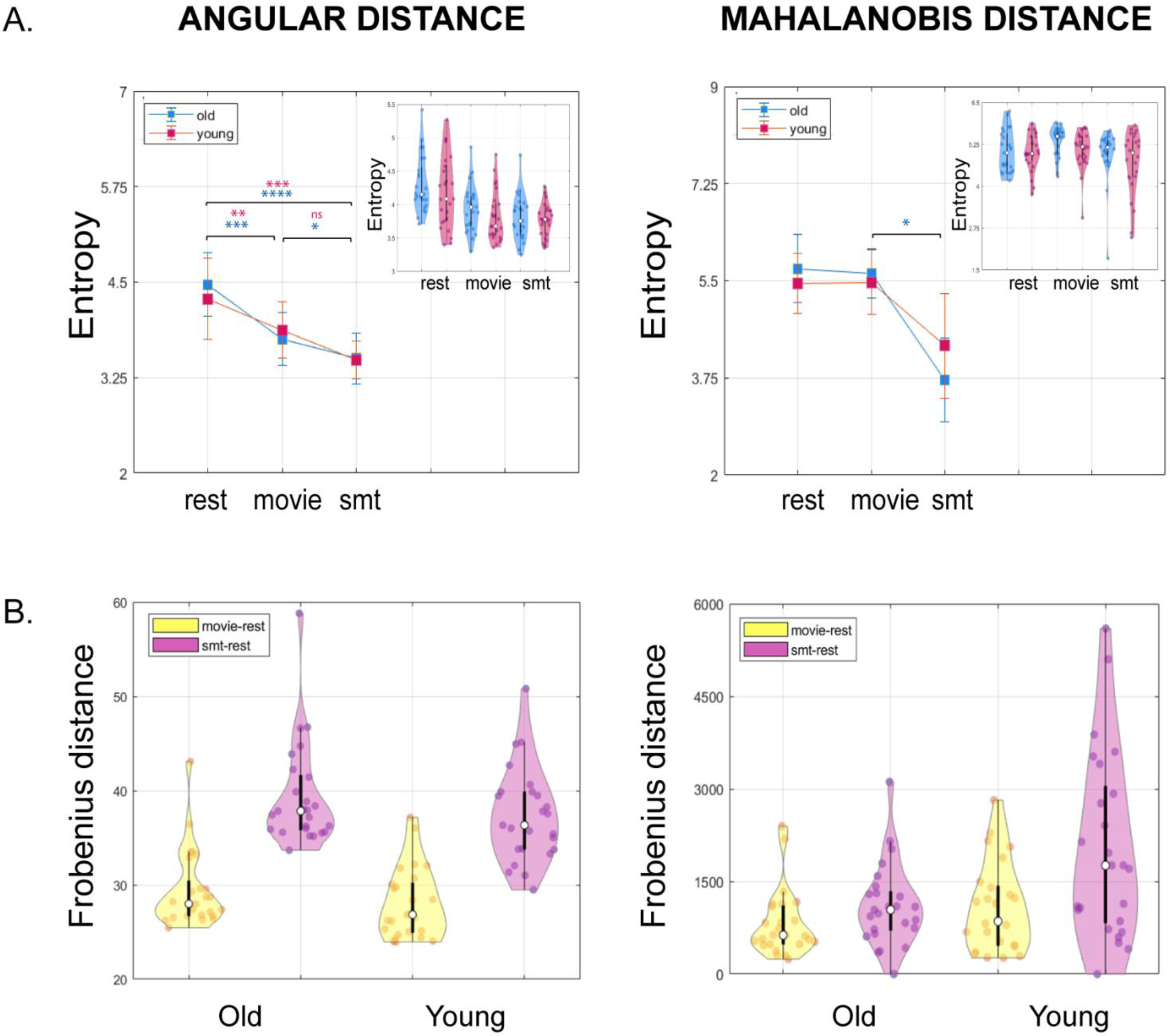
**(A)** Plots representing entropy of temporal stability matrices of resting state (rest), naturalistic movie watching task (movie) and sensorimotor task (SMT), for Angular distance and Mahalanobis distance metric, in both young (magenta) and old adults (blue). Statistically significant differences are indicated using * (𝒫 ≤ 0.05), ** (𝒫 ≤ 0.01), *** (𝒫 ≤ 0.01), *** (𝒫 ≤ 0.001), **** (𝒫 ≤ 0.0001), ns (not significant). **(inset)** Distribution of entropy computed from temporal stability matrices of the resting state, naturalistic movie watching, and sensorimotor task, each contrasted between young (magenta) and old adults (blue) represented as a violin plot **(B)** Plots representing distribution of Frobenius distance between temporal stability matrices of resting state, naturalistic movie watching (yellow) and resting state, sensorimotor task (pink) for Angular distance, and Mahalanobis distance metric, in both young and old adults. The violin plots reveal a shorter Frobenius norm between resting state and movie watching task than resting state and sensorimotor task in both young and old adults.

#### 3.2.2 Using Mahalanobis to quantify temporal stability of *dFC* between young and elderly

Mahalanobis distance between dominant dFC subspaces showed patterns similar to principal angle in young and elderly. Further, we calculate entropy as shown in **Figure 4A**, of temporal stability matrices of each category, in both young and old adults. The results indicate peak entropy in resting state, followed by movie watching task and sensorimotor task, a similar trend as the angular distance metric. In the elderly, the distribution was non-parametric (normality check was done with Jarque-Bera test and D’Agostino-Pearson omnibus test). Wilcoxon matched paired test revealed statistical significance between the entropy of temporal stability matrices of movie watching task and sensorimotor task (p=0.0074, d=0.379). Frobenius norm analysis as shown in **Figure 4B** revealed a shorter Frobenius norm between resting state and movie watching task than resting state and sensorimotor task. The entropy analysis between young and elderly in resting state and tasks is shown in **Figure 4A(inset)**. The analysis indicates entropy of resting state in older adults was higher than their younger counterparts, in both angular distance and Mahalanobis distance metric but statistical tests (independent t-test for angular distance metric and Wilcoxon rank-sum test for Mahalanobis distance metric) did not reveal any statistical significance.

### 3.3 Stochastic characterization of ***dFC***

We examined the stochastic structure of *dFC* evolution by investigating the principal angle *ϕ* (*t*) and Mahalanobis distance *ϕ* (*t*) as functions of time. *ϕ* (*t*) and *ϕ*(*t*) are modelled as auto-regressive or AR (*ρ*) process. The optimal model order was taken to be at the value which yields lowest Akaike information criterion (AIC). The results from this analysis shown in **Figure 5A** and **Figure 5B** reveal the best fit model that explains *ϕ* (*t*) has a model order ρ≥ 6 i.e., the results suggest *ϕ* (*t*) of resting state, movie watching task and sensorimotor task, in both young and old adults, is neither random (ρ≠ 0) nor markovian (ρ≠ 1) in nature, and is dependent on at least 6 immediately preceding values of *ϕ*. For *ϕ*(*t*), as shown in **Figure 5C** and **Figure 5D** both resting state and tasks have the optimum model order *ρ* ≥ 6, suggesting *ϕ*(*t*) is neither random (ρ≠ 0) nor markovian (ρ≠ 1) in both young and old adults.

**Figure 5:**
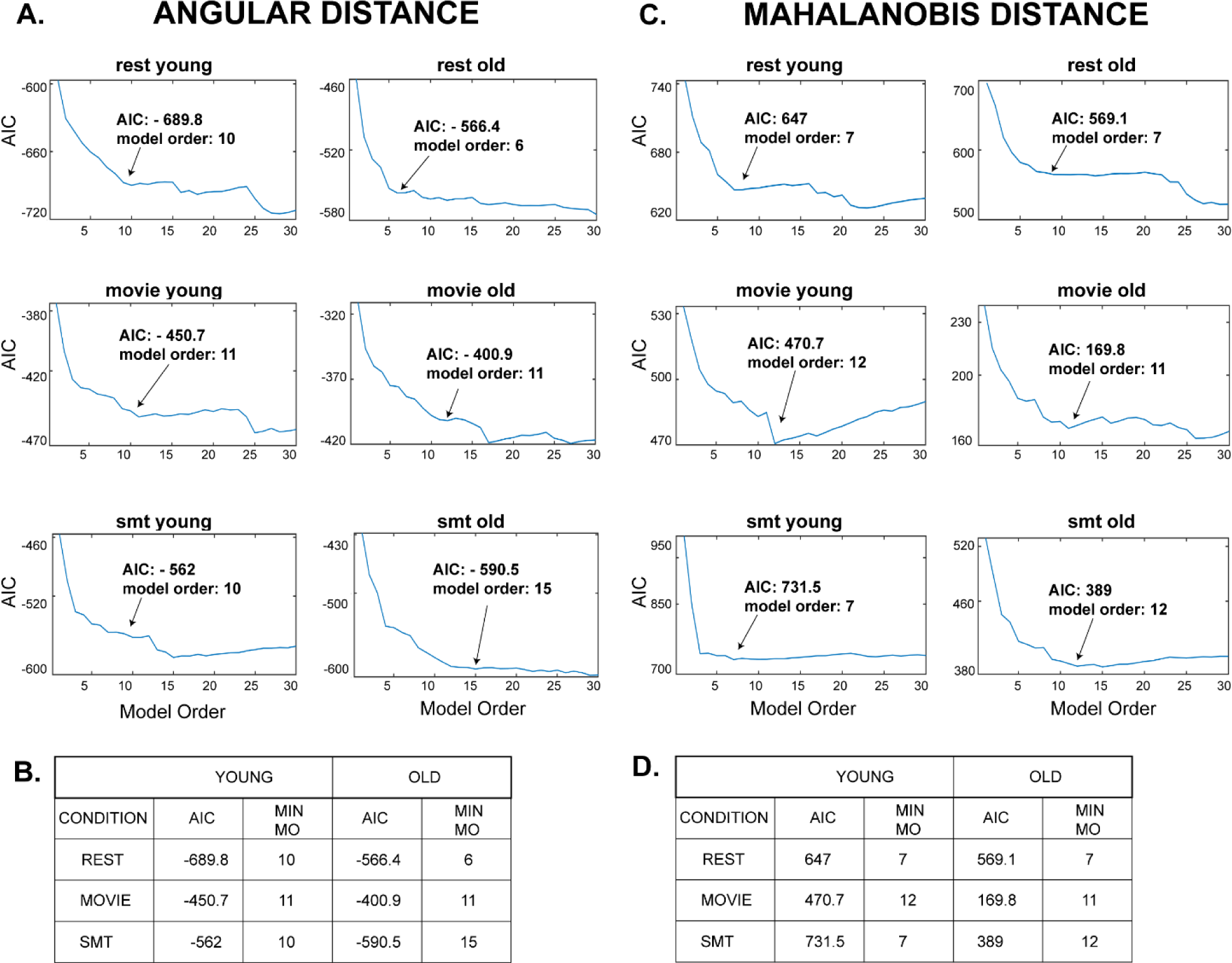
**(A)** Stochastic modelling of principal angle, *ϕ* (*t*) as autoregressive, AR (*ρ*) process. The model order (*ρ*) was varied from 0 to 100. The plot represents Akaike information criterion (AIC) values corresponding to the model order. Inset shows the first minima of the AIC value and its corresponding model order. **(B)** Table shows first minimal AIC value and its corresponding model order of *ϕ* (*t*) for all the categories **(C)** Stochastic modelling of Mahalanobis distance, *ϕ* (*t*) as AR (*ρ*) process. The model order (*ρ*) was varied from 0 to 100. The plot represents AIC values corresponding to the model order. Inset shows the first minima of the AIC value and its corresponding model order. **(D)** Table shows first minimal AIC value and its corresponding model order of *ϕ* (*t*) for all the categories

## 4 Discussion

The functional architecture of the brain is dynamic and changes on a minute temporal scale during resting state and task (Gonzalez-Castillo J., et al., 2015) (Hutchison & et al, 2013) (Gonzalez-Castillo & Bandettini, 2018) (Bolton, Morgenroth, Preti, & Van De Ville, 2020). While previous studies have explored flexibility (Zhang & et al, 2016) (Yin, et al., 2016) and temporal variability (Zhang & et al, 2016) (Li, Lu, & Yan, 2019) of the functional architecture of a specific region, we propose a novel unsupervised method, that captures the stability of whole-brain functional architecture on a minute temporal scale. First, we apply the data-driven unsupervised approach to characterize the high dimensional dynamic functional connectivity into lower dimensional patterns by identifying temporally similar dominant FC configurations. Subsequently, using two different measures - principal angle and Mahalanobis distance applied on dFCs extracted across time, we capture the stability of dFC through the *temporal stability matrices* that could be used to draw critical insights about underlying functional brain states. For empirical validation, we explored modifications in temporal stability matrices of whole-brain FC during a continuous, naturalistic movie watching task and discrete, goal oriented sensorimotor task and showed that, in contrast to resting state, stability increased during the task (stability was highest in the sensorimotor task, followed by naturalistic movie watching task and resting state). Next, we explored ageing specific modulations in temporal stability matrices of dFC patterns between resting state and task and showed increased stability in the task in both young and old adults. Finally, we examined the stochastic properties of temporal stability matrices using an auto-regressive modelling, and showed dominant whole-brain FC configurations are neither random nor Markovian. We discuss the implications of these key results in the following subsections.

### 4.1 Stochastic properties of dynamic functional connectivity

Studies describing brain dynamics have clustered recurring connectivity patterns into states, using clustering algorithms like K-means clustering (Allen, et al., 2014) (Cabral, et al., 2017) (Damaraju, et al., 2014), HMM (Cabral, et al., 2017) (Vidaurre, Smith, & Woolrich, 2017) (Vidaurre, et al., 2016) (Quinn, et al., 2018), suggestive of stability of functional architecture of the brain. Yet, most of the studies hypothesize a fixed number of discrete recurrent connectivity patterns or states with varying temporal fractional occupancy. The homogenous states are essentially clustered ignoring their temporal order and index. Studies have shown clustering time series requires ignoring some data and few attempts at clustering time series have shown to be objectively incorrect in some cases (Rakthanmanon, Keogh, Lonardi, & Evans, 2011) (Rahman, Damaraju, Saha, Plis, & Calhoun, 2020). Rahman and colleagues (Rahman, Damaraju, Saha, Plis, & Calhoun, 2020) have proposed a novel framework, relying on the concept of shapelets, ‘statelets’- a high dimensional state-shape representation of temporal dynamics of functional connectivity, instead of clustering. Another set of prior studies have explored the other side of stability – flexibility, which characterises heterogenous connectivity between a specific region and others over time (Yin, et al., 2016) (Harlalka, Bapi, Vinod, & Roy, 2019) and temporal variability (Zhang & et al, 2016) (Li, Lu, & Yan, 2019) of functional architecture in resting state (Li, Lu, & Yan, 2019) ,naturalistic movie watching task (Li, Lu, & Yan, 2019) and in disease (Zhang & et al, 2016). But these studies are restricted to temporal variability and flexibility of the functional architecture of a specific region. Our main contribution in this study is an unsupervised, data-driven approach to characterise the stability of whole-brain functional connectivity patterns. A recent study (Faghiri, et al., 2020) has proposed a new method, where they calculate the gradients of timeseries pair and use their weighted average of shared trajectory (WAST) as a new estimator of dFC. This method defines a subspace on the raw BOLD fMRI timeseries where as our approach estimated dFC with BOLD phase coherence and defined dominant whole-brain FC patterns as dominant dFC subspaces with PCA and characterised temporally similar dominant whole- brain FC patterns with two alternative measures, angular distance and verifying the same with Mahalanobis distance **(Figure 1B).** The central idea is if the dominant FC configurations are similar for extended time points, then they are considered to be stable.

Viduarre and colleagues (Vidaurre, Smith, & Woolrich, 2017) have shown dynamic switching between brain networks and time spent visiting distinct brain networks are not random. Subsequently, another study has shown that the switching dynamics of functional brain states in the resting state follows AR model of order 1, or in other words a Markovian process fully explains the dFC evolution when correlation was computed using a sliding window approach (liégeois, Laumann, Snyder, Zhou, & Yeo, 2017). By constructing the unsupervised temporal stability matrices from two alternative approaches - principal angle, *ϕ* (*t*) and Mahalanobis distance, *ϕ*(*t*), we reveal that dFC evolution is neither random nor Markovian (**Figure 5A** and **Figure 5B**) (**Figure 5C** and **Figure 5D)**.

### 4.2 Temporal stability of task related dynamic functional connectivity is higher than rest

A key finding of our study indicates a global spread of shorter-lived, repeated patterns of stability between dominant FC configurations in resting state and local spread of longer-lived repeated patterns of stability in the task (in both continuous, naturalistic movie watching task and discrete goal oriented sensorimotor task) (**Figure 2B** and **Figure 3B).** The resting state is shown to be a multistable stationary state-regime at equilibrium (Deco & Jirsa, 2012). Ghosh and colleagues (Ghosh, Rho, McIntosh, Kötter, & Jirsa, 2008) have demonstrated that resting state networks operate close to instability and explore these states, before committing to one of these states. Deco and Jirsa (Deco & Jirsa, 2012) have proposed that a repertoire of multistable states exists in resting state, that are functionally meaningful and inherently supported by the neuroanatomical connectivity, and can be rapidly activated even in the absence of any task. We speculate that in resting state the global spread of shorter-lived repeated patterns of stability between dominant FC configurations is associated with the exploration of multistable dynamic repertoire of states. On the contrary during a task (continuous or discrete), the repertoire of multistable states are limited, as only task specific, cognitively relevant brain networks are explored. The brain visits task specific stable states for duration that a putative stimulus triggered cognitive process demands. This is associated with the local spread of longer-lived temporal similarities between dominant functional connectivity subspaces in a task.

Our entropy results indicate the stability of functional connectivity architecture was highest in the discrete, goal-oriented sensorimotor task, followed by continuous naturalistic movie watching task and resting state **(Figure 4A).** This is in line with previous studies which report an increase in overall stability of FC with the largest increase in between network connections (Elton & Gao, 2015) (Gonzalez-Castillo & Bandettini, 2018), increase in stability of hemispheric homotopic connections during a task (Gonzalez-Castillo J., Hoy, Handwerker, & Bandettini, 2014). Such increased stability of FC during a task is hypothesised to be associated with cognitive constraints during a task (Gonzalez-Castillo & Bandettini, 2018). Frobenius distance analysis results reveal the temporal stability matrices of functional connectivity during continuous, naturalistic movie watching task was closer to resting state than discrete, goal oriented sensorimotor task **(Figure 4B).** Considering our Frobenius distance analysis, we hypothesized stability of functional connectivity architecture should be highest in the sensorimotor task, followed by the naturalistic movie watching task, which was validated by our entropy results. Our findings thus provide evidence of increased temporal stability of whole-brain functional connectivity in task, highest in the discrete, goal-oriented task, followed by continuous, naturalistic movie watching task and then resting-state, using a novel unsupervised approach of characterising the stability of functional connectivity architecture.

### 4.3 Ageing introduces temporal variability in evolution of dynamic functional connectivity in both rest and task

Evidence from prior studies reveals the complexity of FC dynamics remains similar for all participants irrespective of age. An earlier study (Viviano, Raz, Yuan, & Damoiseaux, 2017) found no association between age and rate of switching between the FC states for resting brain. Our results **(Figure 2B** and **Figure 3B)** indicate an overall trend of global spread of shorter-lived repeated patterns of stability between dominant FC configurations in resting state and local spread of longer-lived repeated patterns of stability in the task was similar in both young and old adults. Our study also revealed the highest stability of functional connectivity in the discrete, goal-oriented sensorimotor task, followed by continuous, naturalistic movie watching task and resting state, a trend similar in both young and old adults **(Figure 4A).** Interestingly, McIntosh and colleagues (McIntosh, et al., 2010) have reported BOLD signal variability of hub-region decreases with age, suggestive of increase in stability of hub regions with age. Our results, which contrasted the stability of functional architecture in young and old adults **(Figure 4A (inset)),** found increased stability of functional architecture in young adults in resting state. The neural noise hypothesis suggests the age-related cognitive decline could be explained as a consequence of the increase in the noisy baseline activity of the brain (Voytek, et al., 2015) (Davis, et al., 2009). In accordance to this hypothesis, the decrease in stability of the functional architecture of the brain in older adults can be explained with an increase in neural noise with age. An important point to note, regardless of age associated changes in the stability of functional architecture, our results did not reveal statistically significant differences. Therefore, although there are differences in stability of functional architecture with age, their magnitude may be modest.

### 4.4 Limitations and Future directions

An important caveat of the current study was due to parcellation atlas used in the Cam-CAN dataset. The AAL atlas parcellates the brain regions into 116 structural parcels and few parcels span multiple functional regions. For future studies, for a more refined spatial profile of temporal stability of functional architecture, using a finely parcellated brain atlas is recommended. Researchers have shown stability of functional architecture is modified in patients of Schizophrenia, ADHD and ASD (Zhang & et al, 2016) (Guo, Zhao, Tao, Liu, & Palaniyappan, 2017). Hence, we can extrapolate that the temporal stability of functional architecture can provide a richer information to discover biomarkers for neurological and mental disorders.

### 4.5 Conclusion

In summary, the current study introduces a data-driven unsupervised approach to characterise the temporal stability of functional architecture. When applied to a putative lifespan ageing data, the whole-brain temporal dynamics of naturalistic movie watching task was found to be closer to resting state than during sensorimotor task. Further, the study revealed peak temporal stability in sensorimotor task, followed by naturalistic movie watching task and resting state, a trend similar in both young and elderly. The temporal stability of functional architecture of the resting state was also found to be higher in young adults than their older counterparts. The quantification of differences in network stability associated with healthy ageing provides evidence for the potency of the temporal stability measure to act as biomarker for multiple neurological disorders.

## Acknowledgements

We acknowledge the generous support of NBRC Core funds and the Computing facility. This study was supported by Ramalingaswami Fellowship (Department of Biotechnology, Government of India) to DR (BT/RLF/Re-entry/07/2014) and grant number F.NO. K-15015/42/2018/SP-V from Ministry of Youth Affairs and Sports, Government of India to AB. DR was also supported by SR/CSRI/21/2016 extramural grant from the Department of Science and Technology (DST) Ministry of Science and Technology, Government of India. DR and AB acknowledge the generous support of the NBRC Flagship program BT/ MEDIII/ NBRC/ Flagship/ Program/ 2019: Comparative mapping of common mental disorders (CMD) over lifespan. Data collection and sharing for this project was provided by the Cambridge Centre for Ageing and Neuroscience (CamCAN). CamCAN funding was provided by the UK Biotechnology and Biological Sciences Research Council (grant number BB/H008217/1), together with support from the UK Medical Research Council and University of Cambridge, UK. In accordance with the data usage agreement for CamCAN dataset, the article has been submitted as open access.

## Declaration of competing interest

The authors declare no conflicts of interest

## Ethics statement

CamCAN dataset was collected in compliance with the Helsinki Declaration, and has been approved by the local ethics committee, Cambridgeshire 2 Research Ethics Committee (reference: 10/H0308/50)

## Supplementary Figures

**S 1:**
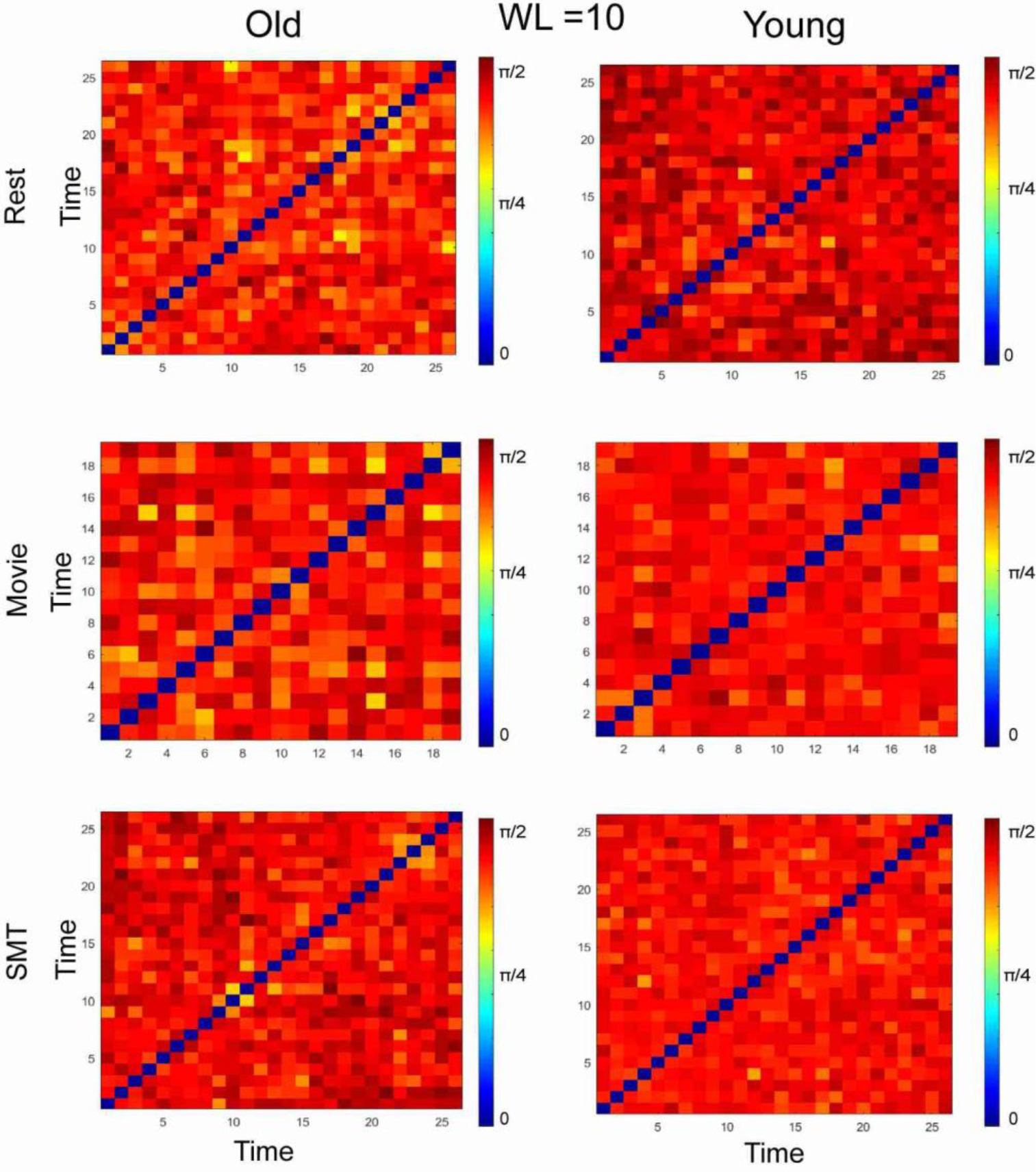
Temporal stability matrices of the resting state, naturalistic movie watching task and sensorimotor task, where each entry is the principal angle *ϕ*(*t*_*x*_, *t*_*y*_) between dominant dFC subspaces at *t*_*x*_ and *t*_*y*_, for young and old adults. For validation of the results where dFC was estimated using BOLD phase coherence, we calculated dFC using sliding window approach with (window length) WL = 10 time points.

**S 2:**
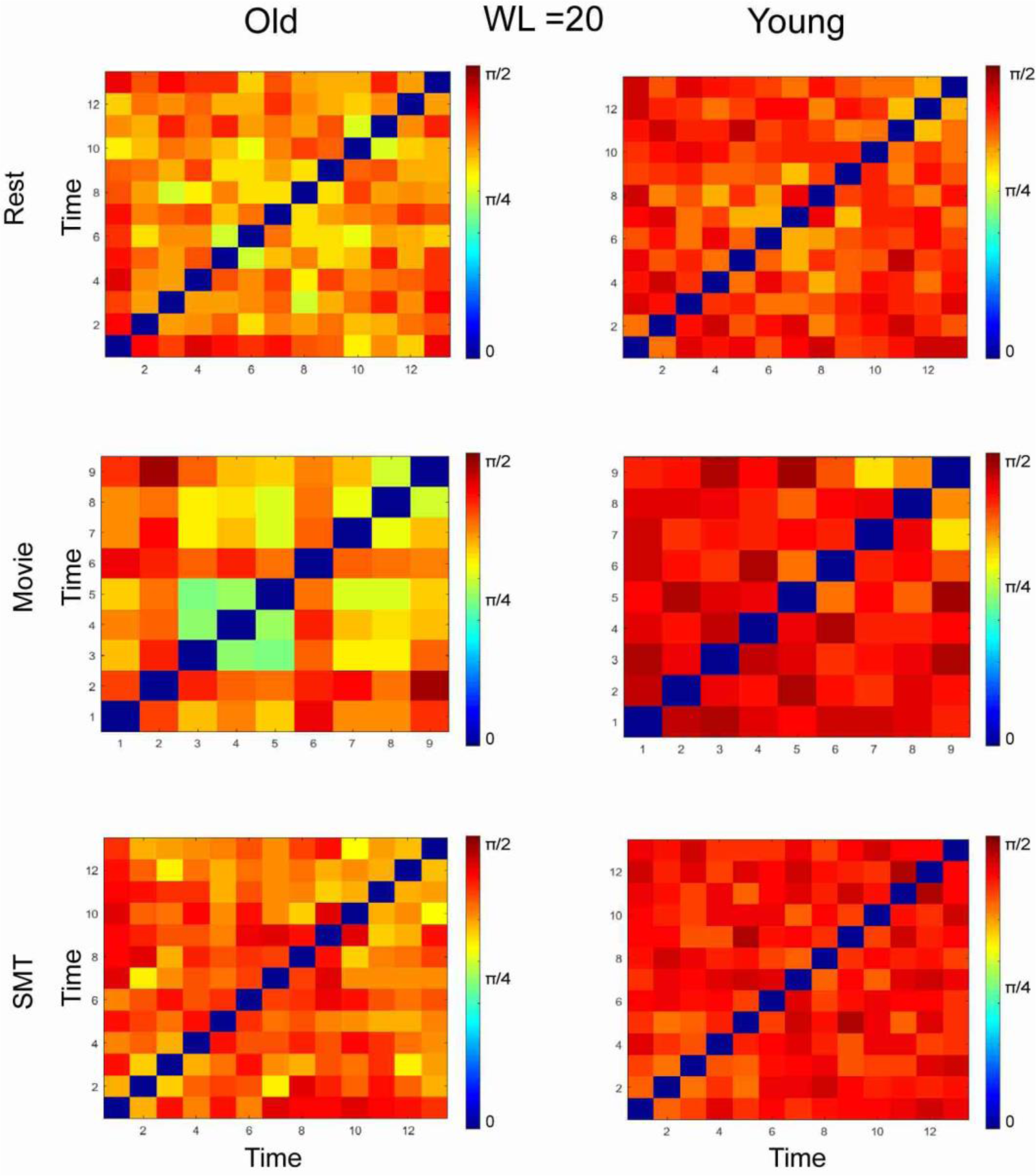
Temporal stability matrices of the resting state, naturalistic movie watching task, and sensorimotor task, for both young and old adults. dFC was estimated using sliding window approach with (window length) WL= 20 time points.

**S 3:**
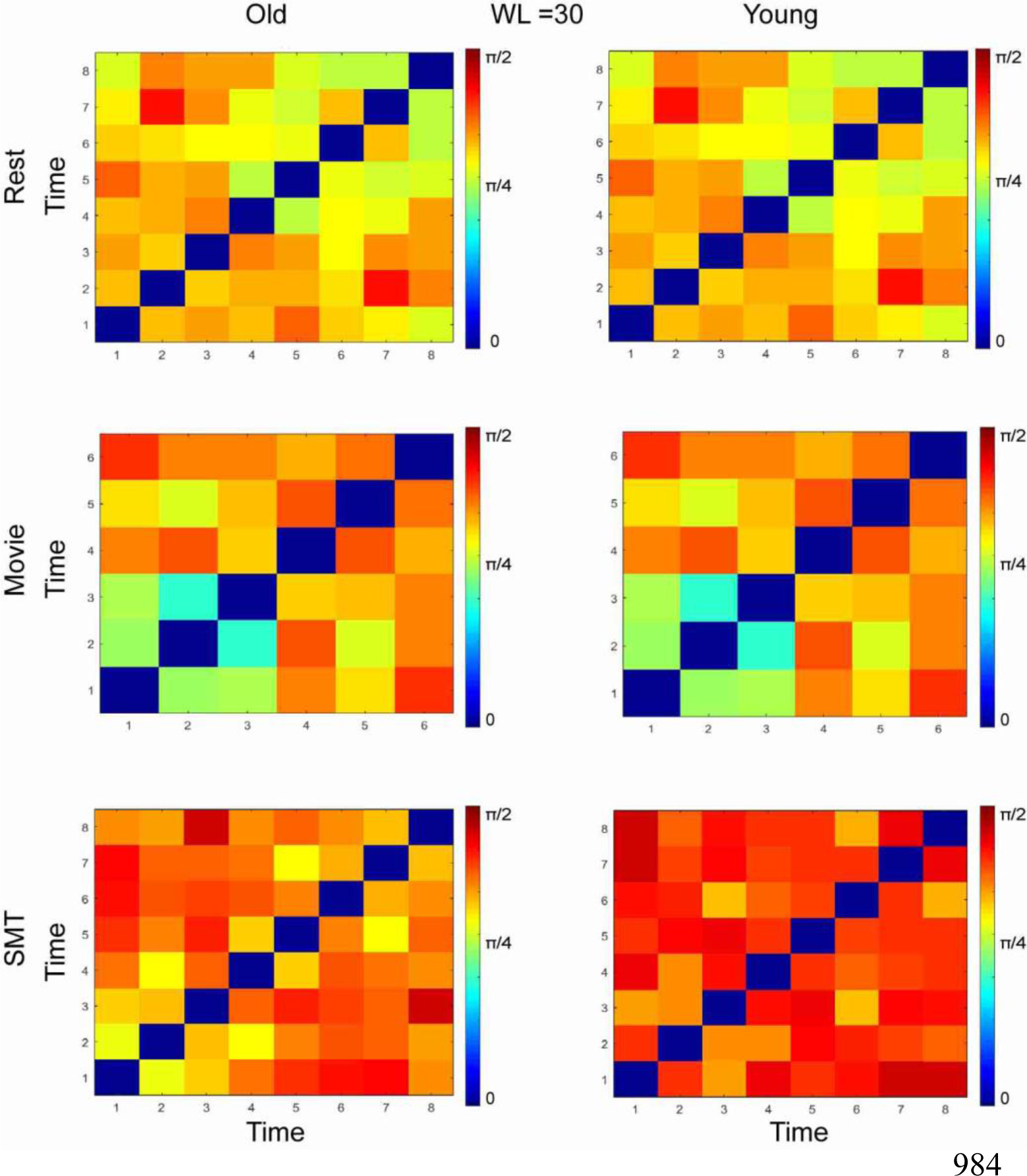
Temporal stability matrices of the resting state, naturalistic movie watching task, and sensorimotor task, for both young and old adults. dFC was estimated using sliding window approach with (window length) WL= 30 time points

**S 4:**
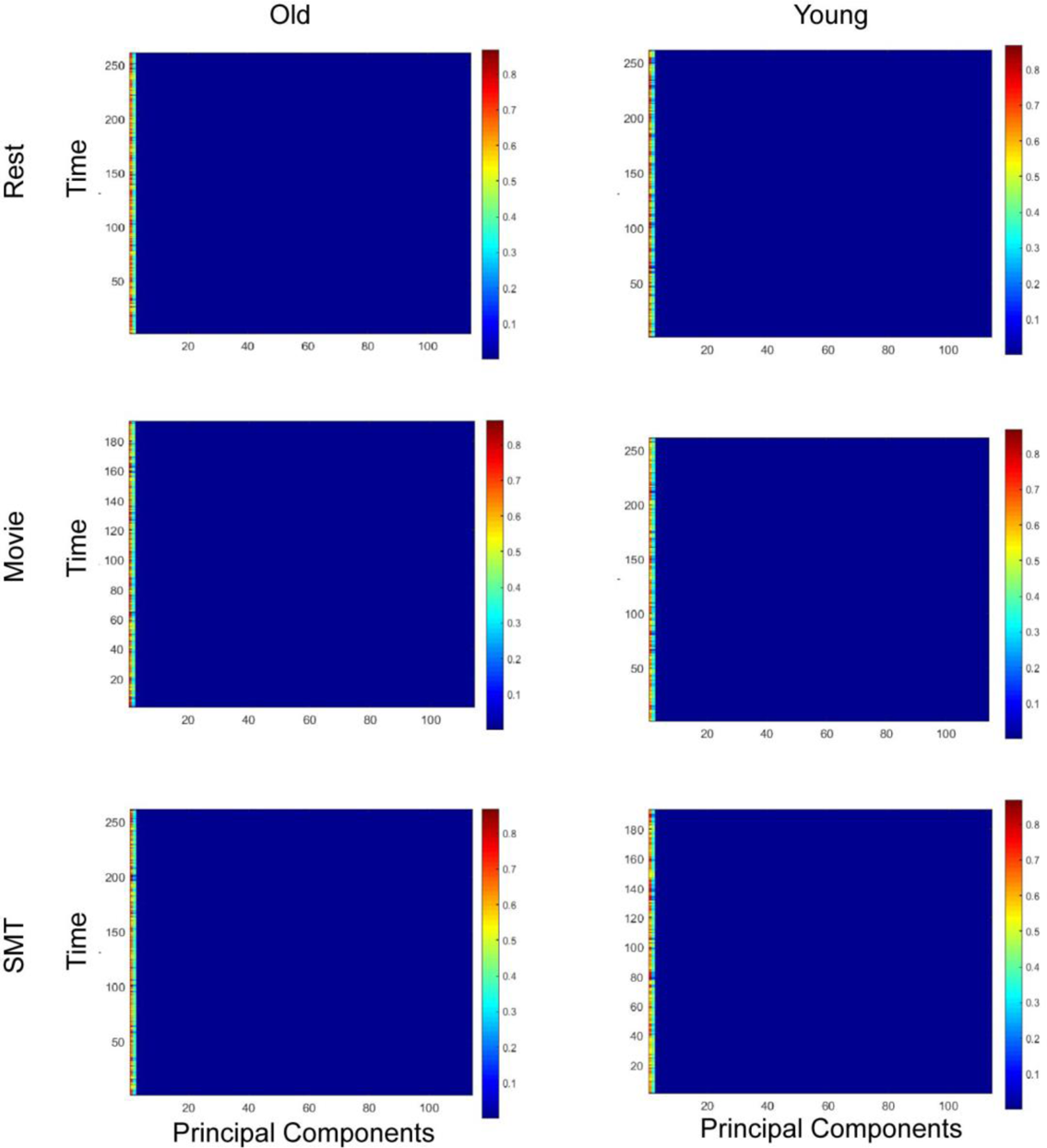
The Plot represents the variance explained by all 116 principal components of the input dFC matrix for all categories. The first three principal components explain almost 99% of the variance of the input matrix.

**S 5:**
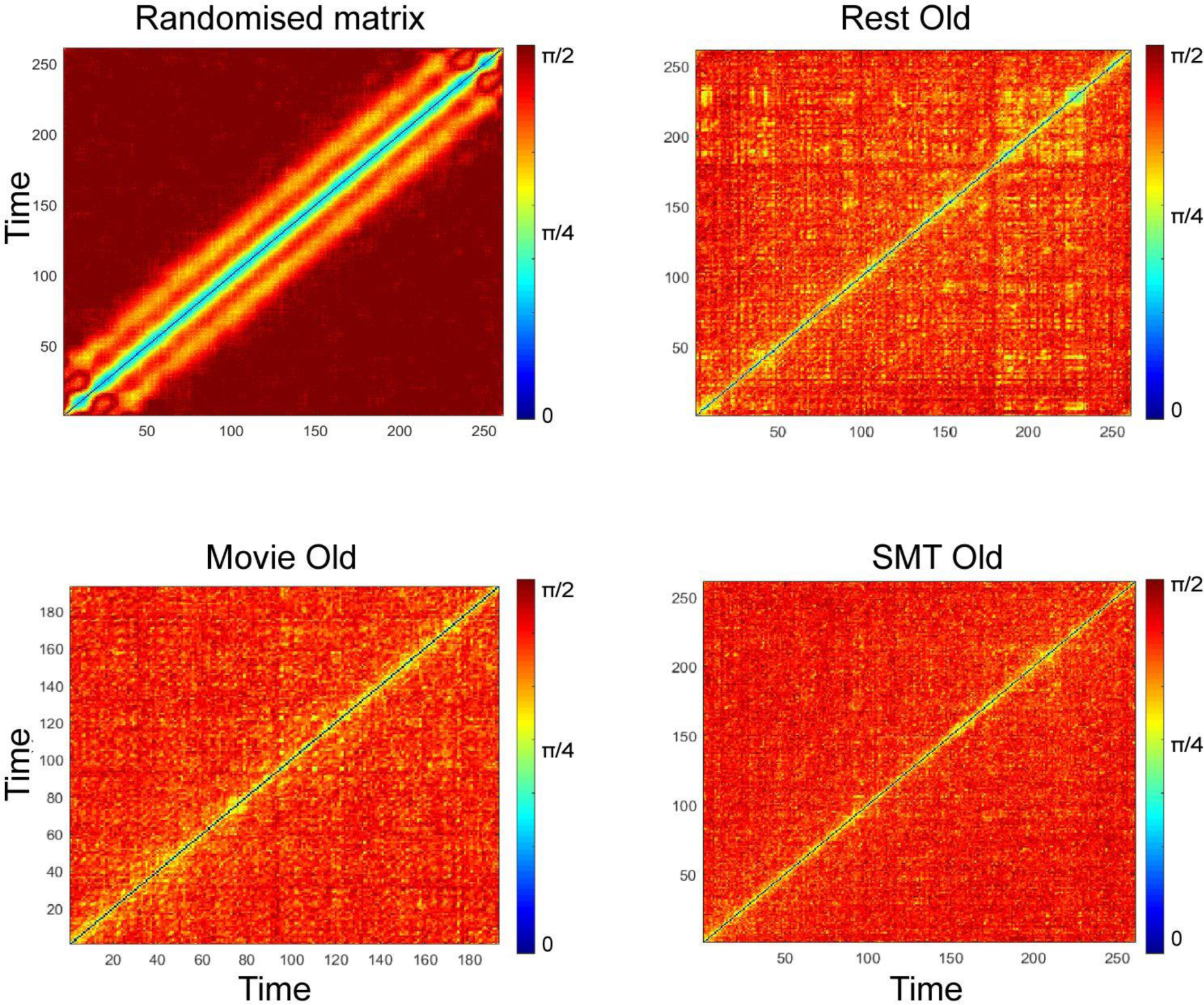
Temporal stability matrices representing the temporal landscape of randomised BOLD signals, resting state, naturalistic movie watching task, and sensorimotor task.

## Notes

### Competing Interest Statement

The authors have declared no competing interest.

